# Benchmarking Free Energy Computational Methods for Revealing the Interactions Driving PARP1 Selective Inhibition

**DOI:** 10.64898/2025.12.29.696816

**Authors:** Alejandro Feito, Natàlia DeMoya-Valenzuela, Cristian Privat, Andrés R. Tejedor, Marco DelValle-Carrillo, Sara Cembellín, Lucía Paniagua-Herranz, Adiran Garaizar, Javier Oller-Iscar, Alberto Ocana, Jorge R. Espinosa

**Affiliations:** Department of Physical Chemistry, Universidad Complutense de Madrid, Av. Complutense s/n, Madrid 28040, Spain; Instituto Pluridisciplinar, Universidad Complutense de Madrid, P.º de Juan XXIII, 1, Moncloa - Aravaca, 28040 Madrid, Spain; Experimental Therapeutics Unit, Hospital Clínico San Carlos (HCSC), Instituto de Investigación Sanitaria San Carlos (IdISSC), Madrid, Spain; Yusuf Hamied Department of Chemistry, University of Cambridge, Lensfield Road, Cambridge CB2 1EW, UK; Department of Organic Chemistry, Universidad Complutense de Madrid, Av. Complutense s/n, Madrid 28040, Spain; Data Science, Bayer AG, Germany; PhAsIca Biosciences S.L, Calle Velázquez, 27, 28001 Madrid, Spain

**Keywords:** PARP inhibitors, molecular dynamics, protein-ligand binding, free-energy calculations

## Abstract

Accurate prediction of inhibitor selectivity across protein paralogues remains a central challenge in computational drug discovery. Here, we systematically benchmark three computational methods—Molecular Mechanics/Poisson–Boltzmann Surface Area (MM/PBSA), free energy perturbation (FEP) and potential of mean force (PMF) calculations—in their ability to recapitulate PARP1 *versus* PARP2 selectivity for eight clinically relevant PARP enzyme inhibitors used in ovarian, breast and prostate tumors among others. We demonstrate how MM/PBSA calculations offer rapid and qualitative insights, but show pronounced sensitivity to the chosen static conformational pose, being particularly challenging for ligands with subtle energetic differences between distinct protein paralogues. In contrast, both FEP and PMF calculations using atomistic models with explicit solvent result in substantially improved agreement with experimental binding affinities. The FEP method exhibits the strongest quantitative correlation with experimental binding free energy differences, remarkably reproducing selectivity trends even among nearly isoenergetic complexes. Notably, our structural contact analysis reveals how contact connectivity controls ligand selectivity, providing valuable mechanistic and molecular insight into the key residues that stabilize each inhibitor in both protein enzymes. Together, our multi-method computational study contributes to elucidate potential chemical modifications across the ligand chemical space to enhance potency and specificity, informing the future design and evaluation of selective inhibitors for precision oncology, including therapies targeting homologous recombination–deficient cancers.

## I. INTRODUCTION

Poly(ADP-ribose) polymerase 1 (PARP1) and 2 (PARP2) are key nuclear enzymes that serve as primary sensors of DNA single-strand breaks and orchestrators of the subsequent repair response^1–3^. Both enzymes bind damaged DNA and catalyze the synthesis of poly(ADP-ribose) chains that drive chromatin relaxation and recruit DNA repair factors^4–7^. Targeting PARP1 and PARP2 with small-molecule inhibitors has become an effective therapeutic strategy^8^, exploiting synthetic lethality in cancer cells with homologous recombination repair deficiency^9,10^. Nevertheless, although PARP1 and PARP2 share overlapping biochemical functions and partially redundant activities^2^, accumulating structural, biochemical, and genetic evidence demonstrates that PARP1 is the dominant mediator of DNA damage detection, whereas PARP2 plays a more specialized role in regulating chromatin organization and hematopoietic cell homeostasis^2,10^. Disrupting PARP1 activity without substantially inhibiting PARP2 is therefore emerging as a powerful strategy for overcoming the hematologic toxicities associated with current PARP inhibitor (PARPi) therapy, in which off-target suppression of PARP2 significantly narrows the therapeutic window^11–13^. This distinction has motivated a new generation of highly PARP1-selective inhibitors—exemplified by Saruparib^14^ and NMS-P118^15^—designed to retain potent antitumor activity while minimizing dose-limiting cytopenias^16^. Therefore, understanding, and more importantly predicting the molecular determinants that generate such selectivity remains a central challenge in structure-guided computational drug discovery^17^.

Protein structure prediction combined with molecular docking provides plausible inhibitor binding modes and positional alignments within enzyme active sites; however, these static representations often overlook conformational plasticity, dynamic water networks, and entropic contributions that critically influence binding specificity^18,19^. Computational free energy calculations can therefore play a central role in elucidating PARP1 *versus* PARP2 selectivity, with complementary strengths and limitations depending on their underlying physical approximations. Among the most widely used approaches, Molecular Mechanics/Poisson–Boltzmann Surface Area (MM/PBSA) calculations^20^ offer rapid endpoint estimates of binding free energies based on molecular mechanics descriptions combined with implicit-solvent models^21^. However, its accuracy partially depends on thorough conformational alignment and approximate entropic treatments. On the other hand, alchemical free energy perturbation (FEP)^22,23^ methods in combination with all-atom force fields enable chemically rigorous predictions by computing absolute binding free energies through thermodynamic transformations between ligand-protein and ligand solvated states. Nevertheless, despite their higher computational cost, its precision can be still challenged by sampling barriers or slow protein rearrangements that accompany ligand binding in dynamic and flexible active sites. Finally, all-atom potential of mean force (PMF)^24,25^ calculations, usually implemented through umbrella sampling or related enhanced-sampling techniques, resolve the free-energy landscape along physically meaningful dissociation coordinates, capturing both enthalpic and entropic contributions as the ligand traverses the conformational landscape of the catalytic cleft^18^. While the binding free energy profile might moderately vary depending on the chosen association/dissociation pathway, the free energy minimum of the bound state is usually rather independent on such pathway^26^. Together, these computational methods, although entailing a certain degree of approximations, provide complementary insights into the thermodynamics and mechanisms of ligand binding and recognition, offering highly valuable molecular interaction insight into the binding affinity between potential drug candidates and protein binding pockets.

In this study, we perform a comprehensive computational benchmark of eight clinically relevant PARP inhibitors—saruparib^14^, NMS-P118^15^, veliparib^27,28^, olaparib^12,28^, rucaparib^29^, niraparib^30^, talazoparib^31^, and pamiparib. Using explicitly solvated atomistic molecular dynamics simulations^32–36^, we characterize the conformational ensembles of each ligand-protein complex and evaluate binding free energies using MM/PBSA^37^, alchemical FEP^38^, and PMF calculations^39,40^. By directly comparing these three methodological approaches across a chemically diverse ligand set, we assess their predictive accuracy, computational performance, sensitivity to structural variability, and ability to capture the energetic determinants underlying protein selectivity. This integrative analysis reveals how subtle differences in the topology, dynamics, and solvation of the different PARP inhibitors translate into distinct binding modes and ultimate protein selectivity. Moreover, our structural contact analysis and energetic decomposition of the involved interactions reveal how contact connectivity controls ligand selectivity, providing mechanistic insight and a quantitative foundation for guiding the rational development of next-generation PARP1-selective therapeutics with improved efficacy and safety profiles.

## COMPUTATIONAL FREE ENERGY METHODS

Binding energetics for all PARP1 and PARP2 complexes have been quantified using three complementary free energy methodologies—MM/PBSA, FEP and PMF calculations—each differing in their associated computational cost and methodological implementation. Their integration as a multi-technique computational platform provides a rigorous and mechanistically coherent thermodynamic description of ligand-protein selectivity.

### Molecular Mechanics/Poisson–Boltzmann Surface Area (MM/PBSA) Calculations

In the MM/PBSA^41^ framework, the binding free energy is estimated from equilibrium ensembles using a molecular-mechanics description of both intramolecular and intermolecular interactions combined with an implicit treatment of solvation. For each representative configuration, the total free energy is expressed as^42^

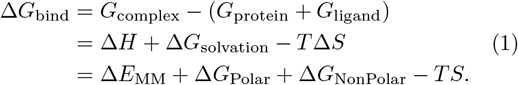

Building on this formalism, the binding free energy is formally defined as the difference between the free energy of the protein–ligand complex and the sum of the free energies of the isolated protein and ligand *G*_complex_ − (*G*_protein_ + *G*_ligand_), which corresponds to the standard thermodynamic cycle underlying all binding free energy calculations. Here, *G*_complex_ denotes the free energy of the bound complex, *G*_protein_ the free energy of the unbound protein, and *G*_ligand_ the free energy of the unbound ligand. This formal definition can be applied to ensembles extracted from extensive MD simulations, but it can also be evaluated using a single representative conformation derived from experimental structures such as X-ray crystallography or cryo-EM.

This definition connects directly to a thermodynamic decomposition in which the free energy change upon binding can be expressed as the sum of an enthalpic term, a solvation contribution, and an entropic penalty respectively (Δ*H* +Δ*G*_solvation_ −*T* Δ*S*), linking the macroscopic thermodynamic description of molecular recognition with the microscopic energetic and entropic processes involved in ligand binding. Here, Δ*H* corresponds to the enthalpic change in intramolecular and intermolecular interactions upon binding, Δ*G*_solvation_ accounts for desolvation effects, and *T* Δ*S* represents the entropic cost of constraining the relative motion and orientation of the binding molecule.

MM/PBSA provides a practical computational realization of this thermodynamic picture by rewriting these contributions in terms of quantities obtained from molecular mechanics and continuum solvation models: (i) the enthalpic component is approximated by the molecular mechanics energy (Δ*E*_MM_), which includes bonded, electrostatic, and van der Waals interactions; (ii) the solvation term is separated into a polar contribution (Δ*G*_Polar_), computed using either Poisson-Boltzmann (PB) or generalized Born (GB) implicit-solvent models in MM/GBSA^43,44^ calculations, and a non-polar component (Δ*G*_NonPolar_), estimated from the solvent-accessible surface area (SASA) and associated with hydrophobic desolvation; and (iii) the entropic term (*S*), which accounts for the configurational entropy loss upon binding, typically evaluated through quasi-harmonic (QHA) or normal-mode analysis (NMODE).

While both the Poisson–Boltzmann (PB) and the generalized-Born (GB) approaches seek to approximate the electrostatic contribution to solvation, they differ fundamentally in mathematical rigor and computational cost. The PB model solves the Poisson–Boltzmann equation numerically^45^,

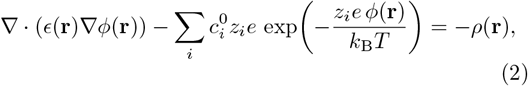

where it describes how the electrostatic potential *ϕ*(**r**) is distributed around a biomolecule in an ionic solution. In this framework, *ϵ*(**r**) denotes the spatially dependent dielectric constant, which takes low values inside the protein cavity and high values in the solvent. The term *ρ*(**r**) represents the fixed charge density of the solute, typically arising from atomic partial charges. The summation term compensates for the contribution of mobile ions in solution, where 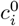 represents the bulk concentration of ionic species *i, z*_*i*_ is their valence, and *e* is the elementary charge. The exponential factor reflects the Boltzmann distribution of ions in response to the local electrostatic potential, with *k*_B_ the Boltzmann constant and *T* the temperature. In the linearized form of the equation, the ionic response is approximated by the Debye–Hückel length *κ*, which encodes the screening effect of electrolytes in solution. Altogether, the Poisson– Boltzmann equation provides a continuum description of electrostatic interactions by coupling the fixed charges of the solute with the redistributable charges of the surrounding ionic environment, providing a more accurate continuum description of the electrostatic environment at the cost of a significantly higher computational demand. In contrast, GB models—including the modern GB5^46^ variant—approximate the PB solution analytically through pairwise de-screening functions, achieving a substantial reduction in computational cost while maintaining reasonable accuracy for most biomolecular systems. GB5 in particular improves the effective Born radius calculation and dielectric screening terms, yielding results that more closely approach PB-level accuracy but still fall short in highly charged or topologically complex environments. In practice, PB is preferred when accuracy is critical^47^, whereas GB5 offers a practical balance between speed and precision for large-scale or highthroughput MM/GBSA calculations. In Section SI of the Supplementary Material (SM), we provide further details of these two methodologies.

### Alchemical Free Energy Perturbation (FEP) Calculations

To obtain chemically accurate estimates of protein-ligand binding affinities, we perform atomistic alchemical FEP simulations in which we gradually transform ligand *A* into ligand *B* using a coupling parameter *λ* that interpolates between the two Hamiltonians. The associated free energy change is computed using the Zwanzig relation^48^:

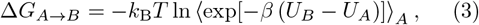

where *U*_*A*_ and *U*_*B*_ refer to the potential energies of the system under the Hamiltonians of states *A* and *B*, respectively; *k*_B_ is Boltzmann’s constant; *T* is the absolute temperature; *β* = (*k*_B_*T*)^*−*1^; and ⟨· · · ⟩_*A*_ denotes an ensemble average over configurations sampled in both states *A* and *B*. This relation provides a rigorous statistical-mechanical framework to compute free energy differences between two chemical states from an ensemble of microscopic configurations.

To ensure thermodynamic consistency, positional restraints need to be applied during the alchemical transformations. Harmonic restraints prevent translation of the ligand within the binding pocket, while additional restraints suppress global rotation and translation of the protein. Specifically, ligand translation needs to be restricted when varying the Hamiltonian by applying the positional restraints to a single heavy atom, minimizing perturbation to the rest of the molecule. In the protein, four strategically selected residues are restrained to suppress overall translation and rotation. This approach preserves a well-defined relative frame of reference between the protein and ligand while allowing the system sufficient structural flexibility to undergo conformational changes. Such restraints ensure stable overlap of conformational ensembles across intermediate *λ* states connecting both Hamiltonians, *A* and *B*, where molecular interactions are partially switched off and positional alignment cannot be maintained through the inherent intermolecular interactions of the system.

In the alchemical route, the gradual decoupling of the ligand from its environment follows the standard two-stage scheme employed in absolute binding free energy calculations^49^. First, the electrostatic interactions are turned off smoothly as *λ* increases, thereby neutralizing the ligand without introducing large steric perturbations. Once the charges have been fully removed, the van der Waals interactions are gradually scaled down using soft-core potentials that prevent numerical instabilities of atoms losing their excluded-volume repulsion. This sequential treatment—electrostatics first, van der Waals second—ensures a physically meaningful pathway with improved configuration overlap between neighboring windows^50^.

Free energy differences along the full *λ* schedule are subsequently combined using the multi-state Bennett acceptance ratio (MBAR)^51^, which provides a statistically optimal estimator of Δ*G* by explicitly exploiting the overlap of configurational ensembles between adjacent *λ*-windows. In essence, MBAR generalizes the Bennett acceptance ratio and can be expressed as^51^

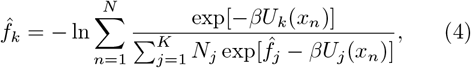

Where 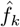 accounts for the dimensionless free energy of state *k, U*_*k*_(*x*_*n*_) is the potential energy of configuration *x*_*n*_ evaluated in the state *k, N*_*j*_ is the number of samples collected in state *j*, and the sum runs over all collected configurations across *K* states. This formulation effectively performs an overlap sampling^52^ by reweighting contributions from all windows according to how well each configuration represents the neighboring states, ensuring robust and statistically efficient estimates of Δ*G*. Relative binding free energies, Δ*G*_bind_, can be finally obtained by subtracting the corresponding alchemical transformation in aqueous solution, thereby connecting the free energy difference in the bound state with that in the unbound solvated state. In this way, Eq. 3 represents the formal statistical-mechanical definition of the alchemical free energy change, while the *λ*-dependent simulations and the corresponding MBAR analysis provide the practical computational implementation used to compute binding free energies from molecular simulations. Please see Section SII-III of the SM for further details of the system preparation and method implementation in this work, respectively.

### Potential of mean force (PMF) calculations

Finally, to complement both the end-point and alchemical FEP calculations, PMF simulations are employed to resolve the free energy landscape of ligand binding association along a physically meaningful reaction coordinate. The ligand was incrementally displaced from the catalytic binding site of PARP1 (or PARP2) to the bulk solution, respectively, while applying harmonic biasing potentials in each umbrella window^39^:

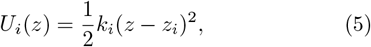

where *z* represents the ligand–protein separation reaction coordinate, *z*_*i*_ is the window center, and *k*_*i*_ is the force constant. Because PMF reconstruction is sensitive to global rigid-body motions, both the protein and the ligand are subjected to partial rotational and translational restraints to ensure a stable reference frame throughout the dissociation/association pathway. In the ligand, these restraints are applied to approximately six heavy atoms, limiting overall translation while allowing internal flexibility and partial conformational rearrangements. In the protein, four selected residues are restrained to suppress global rotation and translation. These restraints prevent spurious drift and ensure that the reaction coordinate reflects genuine unbinding events rather than system-level displacements, while still allowing the system to explore relevant conformational changes.

The unbiased potential of mean force *A*(*s*) is calculated by combining the probability distributions from all umbrella sampling windows and correcting for the biasing potential *w*(*s*) applied in each window, which can be expressed as^53^:

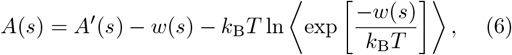

where *A*^*′*^(*s*) is the PMF from the biased simulation, *w*(*s*) is the harmonic biasing potential, *k*_B_ is the Boltzmann constant, *T* is the temperature, and the angle brackets denote an average over the simulation windows. The final, unbiased PMF *W* (*z*) is then reconstructed from all windows using the weighted histogram analysis method (WHAM), yielding a continuous free energy profile that resolves the interplay of enthalpic interactions, conformational flexibility, hydration, and entropic penalties associated with confinement in the catalytic binding pocket. After applying standard-state corrections, the depth of the PMF yields an absolute estimate of the binding free energy (Δ*G*_bind_). Please see Section SII-IV of the SM for further details of the system preparation and implemented methodology in this work, respectively.

Together, these three complementary methodological layers—MM/PBSA evaluation, alchemical FEP transformations analyzed with MBAR, and PMF-based reconstruction across the dissociation pathway—provide a coherent, complementary and quantitatively grounded framework for dissecting the molecular origins of selective PARP1 inhibition. Through this multi-step methodology, we evaluate the relative binding free energy differences (ΔΔ*G*_bind_) between PARP1 and PARP2 for each ligand, enabling a systematic assessment of ligand selectivity across these homologous protein targets.

In Figure 1 we provide a schematic overview of the computational strategies employed to estimate protein–ligand binding free energies. The workflow begins with the preparation of the initial protein–ligand complex configuration, which is subjected to atomistic molecular dynamics (MD) simulations in explicit solvent, including water molecules and ions. Using GROMACS (version 2023)^54^, all simulations are performed with the atomistic a99SB-*disp* force field^55^ for the protein-protein interactions, the TIP4P-*disp* water model, and ligand parameters derived from the OpenFF^56^ toolkit, ensuring a consistent and accurate description of protein, ligand and water/ion interactions. Both PMF and FEP calculations are performed using this force field. In contrast, MM/PBSA can be performed directly on the initial structures experimentally resolved—the following PDB codes are used in this study; PARP1: 9ETQ^14^, 7KK4^28^, 7KK6^28^, 5A00^15^, 7KK5^28^, 6VKK^57^, 7CMW^58^, 7KK3^28^; PARP2: 4TVJ^28^, 3KJD^27^, 4ZZY^15^, 8HLQ^59^, 8HKO^59^, 8HKS^59^, 4PJV^60^—providing a computationally quick end-point estimation of binding affinity. MM/PBSA decomposes the total energy into molecular mechanics and solvation contributions, using the Poisson–Boltzmann equation for polar solvation. FEP employs a gradual alchemical decoupling of the ligand, separating van der Waals and electrostatic interactions across a series of *λ*-windows, with the resulting free energy differences combined via multistate estimators such as MBAR, capturing subtle enthalpic, entropic and solvation effects with minimal positional restraints. Finally, PMF simulations compute the free-energy profile along a predefined reaction coordinate, typically by pulling the ligand out of the binding pocket with controlled positional restraints, thereby characterizing the energetic barriers and intermediate states along the unbinding pathway. The schematic from Fig. 1 also highlights the relative computational cost of the three methods, with MM/PBSA being the fastest, followed by FEP and PMF, and shows how all approaches ultimately converge on the calculation of the binding free energy, Δ*G*_bind_, enabling a comparative assessment of ligand binding affinities and selectivity across homologous protein targets.

**FIG. 1.**
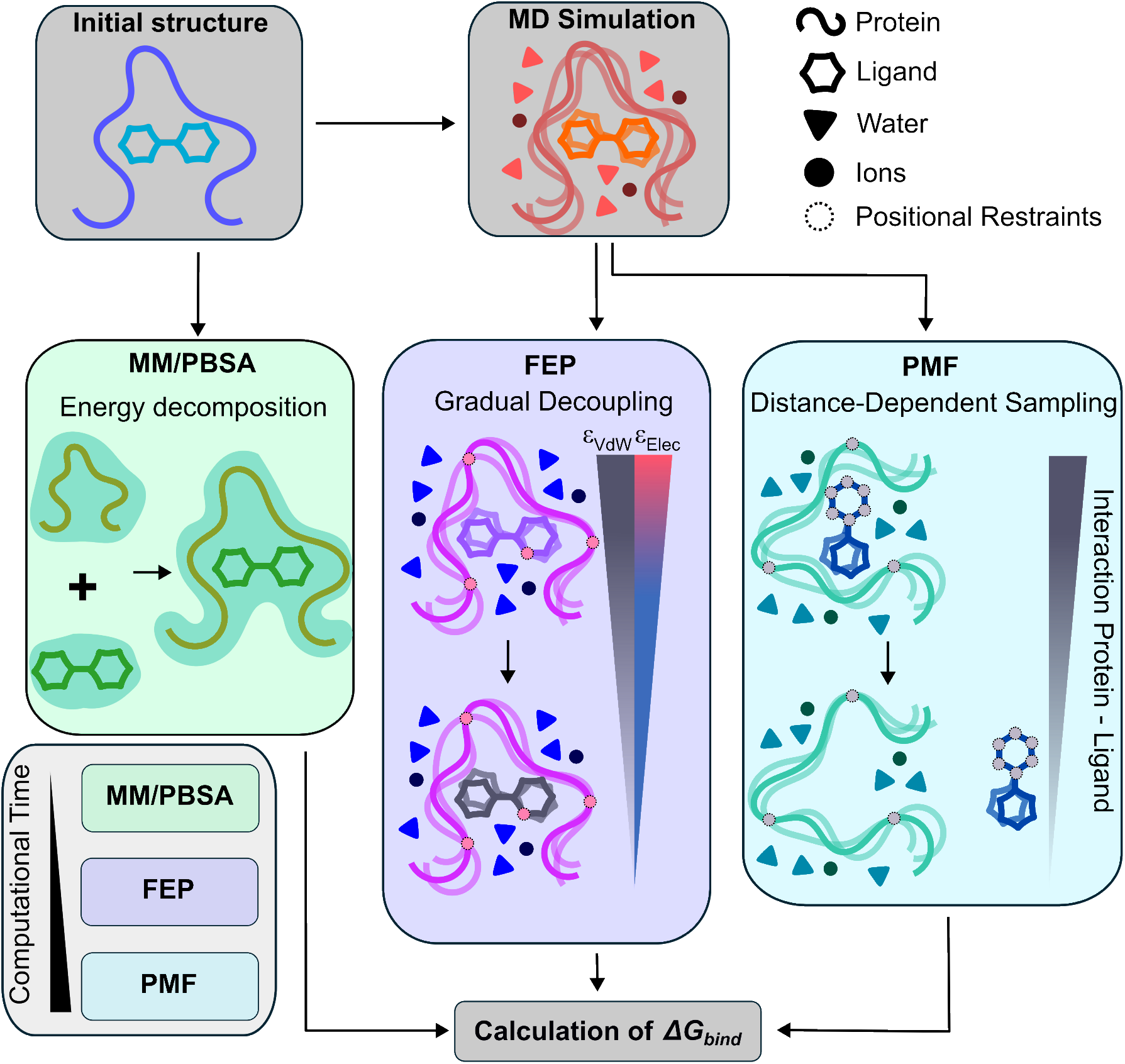
Schematic representation of the three computational approaches used in this work to estimate the binding free energy (Δ*G*_*bind*_) of a given ligand in different protein paralogues. The relative binding affinity can be evaluated by: MM/PBSA, which estimates the free energy through energy decomposition of representative conformations; Free Energy Perturbation (FEP), where interactions between the ligand and its environment are gradually decoupled; and Potential Mean Force (PMF), which quantifies the free energy profile as a function of the protein–ligand dissociation pathway. The relative computational cost of each method is also included.

## RESULTS AND DISCUSSION

### Computational Assessment of PARP1 vs PARP2 Inhibitor Selectivity

Benchmarking computational methods for predicting PARP inhibitor selectivity is essential for assessing the reliability of free energy calculations that can guide the design of more potent inhibitors in drug discovery. Our goal in this section is therefore twofold: first, to examine how the predicted PARP1/PARP2 selectivity evolves as increasingly rigorous and time-demanding computational methodologies are applied; and second, to assess whether these distinct approaches—MM/PBSA, FEP, and PMF—converge on consistent qualitative trends across a chemically diverse set of inhibitors. To this end, we first focus on four representative ligands: saruparib and NMS-P118, which are well-characterized as highly selective PARP1 inhibitors^14,15^, and veliparib and olaparib, which are known to exhibit broader, non-selective activity within the PARP protein family^27,28^. All simulations are performed exclusively on the catalytic binding site of each protein with the ligands, see Section SIV of the SM for further details of the PDB structures and PARP1/2 active site sequence. By including ligands that span these two extremes of selectivity, we ensure that the comparative analysis challenges each computational method not only with subtle energetic differences but also with cases where substantial shifts in binding preference take place. This framework allows us to evaluate not only the numerical accuracy of each technique, but also the stability and robustness of their relative predictions as simulation complexity increases.

To establish the context for this comparison, Figure 2A shows the approximate CPU time required to perform these calculations on a single CPU core with these three different methodologies. A striking feature of this panel is the difference in scale: MM/PBSA differs by several orders of magnitude in comparison with FEP and PMF methods. Such major difference arise naturally from the distinct level of approximations on which each method is built. MM/PBSA relies primarily on static energetic evaluations with an implicit solvent representation, whereas FEP and PMF achieve greater thermodynamic rigor through explicit sampling of protein–ligand interactions and explicit solvent dynamics. As a result, direct comparison of absolute binding free energies across methods is neither meaningful nor expected; instead, the emphasis must be placed on relative patterns between distinct inhibitor binding free energies. The selectivity is thus quantified through the relative binding free energy difference, ΔΔ*G*_bind_, defined as

**FIG. 2.**
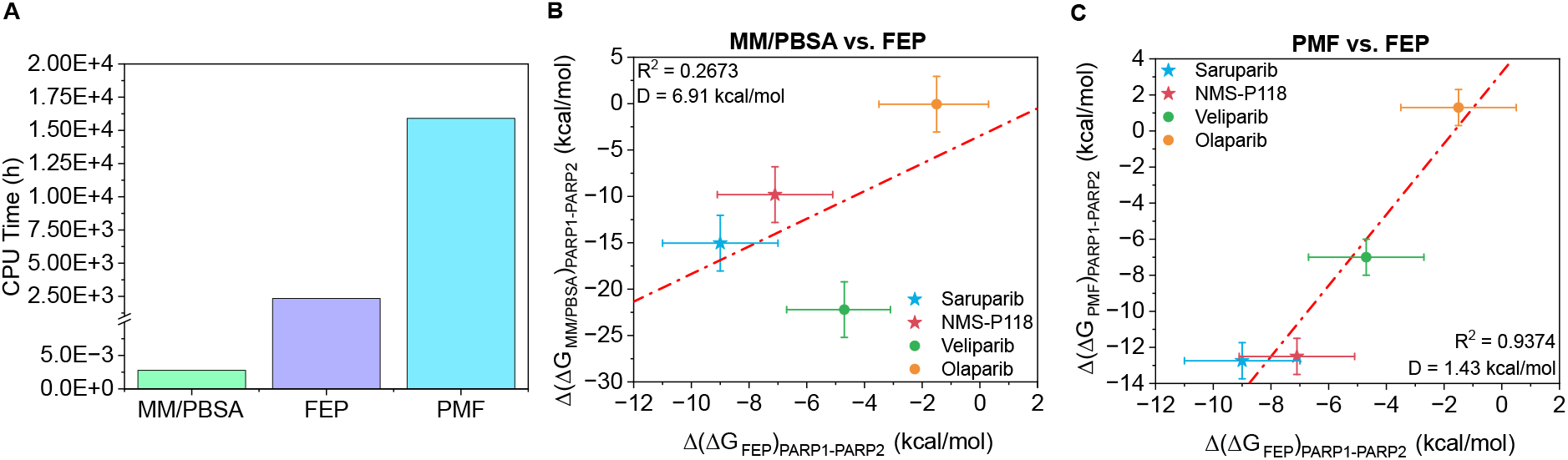
(A) Required simulation time (in hours) of a single CPU core for performing the binding energy prediction using the different tested methods. Comparison of the difference (Δ) of binding free energy (Δ*G*) between PARP1 and PARP2 (Δ(Δ*G*)_*PARP* 1*−PARP* 2_) for four inhibitors: saruparib (blue star), NMS-P118 (red star), veliparib (green circle) and olaparib (yellow circle) using different computational approaches. (B) Correlation between MM/PBSA and FEP results. (C) Correlation between PMF and FEP calculations. Error bars represent the standard deviation of the simulations. Specific inhibitors of PARP1 are plotted as stars and non-specific inhibitors as circles. The red dashed line depicts the linear regression of the data shown.

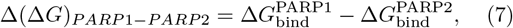

where 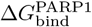 and 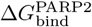 denote the binding free energy of the same ligand to PARP1 and PARP2, respectively.

This perspective becomes clearer in Figure 2B, where we initially compare MM/PBSA and FEP predictions. Despite their methodological differences, the two approaches reveal a consistent linear trend for three of the studied inhibitors: saruparib, NMS-P118, and olaparib. The correlation for these compounds indicates that, although the absolute values differ substantially between MM/PBSA and FEP, both methods capture similar relative selectivity relationships. However, the overall coefficient of determination is modest, with *R*^2^ = 0.2673, reflecting that the linear agreement is limited when considering all four ligands together. Veliparib is chiefly responsible for this reduction in the correlation, as it does not follow the trend defined by the other inhibitors and appears as a clear offset. This deviation strongly suggests that MM/PBSA cannot adequately account for subtle structural and solvation features of veliparib with respect to the rest of the ligands, which are better rep-resented in atomistic FEP simulations. The issue likely stems not from an intrinsic anomaly of the ligand, but rather from limitations of the MM/PBSA protocol— particularly its sensitivity to subtle conformational rearrangements of the molecule within the binding site and solvent-mediated intermolecular interactions. Veliparib thus serves as a diagnostic case illustrating where increased sampling rigor and detailed atomistic interactions begin to substantially influence the predicted selectivity.

To quantify how much MM/PBSA *versus* FEP predictions deviate from an ideal perfect linear relationship, we compute a deviation metric *D*, defined as:

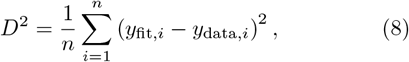

where *n* is the number of data points (ligands), *y*_fit,*i*_ is the value on the regression line (the ideal fitted value), and *y*_data,*i*_ is the actual computed simulation value. In essence, *D* measures the mean squared deviation of the data from the ideal linear fit, where smaller *D* values indicate that the points cluster more tightly around the correlation line.

In our comparison of MM/PBSA *versus* FEP calculations, we obtain *D* = 6.91 kcal/mol, reflecting a considerable deviation of veliparib’s prediction around the fitted line. In the context of the intrinsic uncertainties of each method: MM/PBSA itself is known to exhibit relatively large errors, often around 2–3 kcal/mol^61^, while FEP is generally slightly more precise, with typical uncertainties of 1–2 kcal/mol^62^. Therefore, part of the scatter and the modest *R*^2^ value observed in Figure 2B can be attributed to the intrinsic limitations of both methods rather than to their associated statistical uncertainty.

In Figure 2C, we compare FEP and PMF calculations, two advanced methodologies that entail all-atom MD simulations with explicit solvent. Here, the agreement is markedly stronger: all four inhibitors lie close to a linear trend, with a high coefficient of determination *R*^2^ = 0.9374. This strong correlation suggests that FEP and PMF largely converge on a consistent description of PARP1 *versus* PARP2 selectivity for the considered PARP inhibitors. For FEP *versus* PMF comparison, the deviation metric is *D* = 1.43 kcal/mol, indicating a much closer alignment between these two methods than between MM/PBSA with FEP. The small *D* is consistent with the typical uncertainties of both methods, where PMF calculations usually entail statistical uncertainties of ∼1 kcal/mol^63^. Thus, the observed spread is comparable to the intrinsic error of the methods, and most of the residual scatter likely reflects statistical fluctuations rather than systematic disagreement.

Although FEP and PMF simulations show strong overall agreement, the two approaches moderately differ in the level of discrimination they provide among the considered inhibitors. In particular, while FEP captures noticeable differences across all four compounds, the PMF profiles for the two selective inhibitors (see Fig. SI in the SM) yielded nearly identical free energy values. This behavior reflects several methodological limitations that affect each technique in distinct ways. Two important limitations of FEP are directly related to the statistical efficiency of the alchemical sampling. First, FEP requires sufficient phase-space overlap between adjacent *λ*-windows. When the configurational ensembles at successive *λ* values become too dissimilar, the overlap of energy distributions deteriorates, leading to large variances in the exponential averaging estimator and poor simulation convergence. This phenomenon has been well documented as a major source of noise and instability in FEP calculations^64^. Second, there is a non-trivial interplay between simulation time and statistical efficiency. Although longer simulations per window can, in principle, improve sampling, they may also exacerbate divergence between windows if slow conformational rearrangements occur during the alchemical transformation. As a result, excessive per-window simulation time can counterintuitively reduce phase-space overlap and degrade the accuracy of the free energy estimates. This issue has been highlighted in recent methodological analyses, which recommend using multiple independent windows with moderate sampling rather than excessively long trajectories in each window^65,66^. For these reasons, in our study the FEP protocol is constructed using the same number of *λ*-windows for all four inhibitor–protein systems, and the same number of restraints in each system. This uniform setup ensures that any differences in the resulting free energy estimates arise from the physical behavior of the atomistic force field rather than from methodological inconsistencies in window spacing or statistical sampling. By standardizing the alchemical resolution across all transformations, we minimize the possibility that variations in phase-space overlap or sampling quality can introduce artificial discrepancies in the comparison between different PARP inhibitors.

In contrast, PMF calculations intrinsically rely on a predefined reaction coordinate^67,68^ along which the ligand is pulled out of the binding pocket. The resulting free energy profile is therefore sensitive to the choice of the coordinate: if it does not reflect the physically relevant unbinding pathway, or if multiple pathways exist but are not sampled, the PMF may underestimate or oversmooth energetic differences between ligands. Moreover, PMF calculations typically require positional restraints on the ligand to maintain a controlled pulling trajectory (see Section SIV for further methodological details on the PMF calculations and obtained PMF profiles). These restraints can hinder molecular rearrangements and lead to artificially overestimated free energies barriers across the dissociation pathway^69^. For this reason, in our study the same reaction coordinate and the same number of restrictions are used for all four inhibitors, ensuring that differences between PMF profiles do not arise from arbitrary choices across the pulling path. Nevertheless, it must be acknowledged that any fixed reaction coordinate implicitly favors certain unbinding pathways over others, and may therefore advantage some ligand–protein complexes relative to others in an indirect subtle manner, which is extremely difficult to quantify.

Altogether, these considerations reveal a coherent progression: while MM/PBSA can provide a rapid, approximate qualitative picture, the more rigorous FEP and PMF methods progressively reduce uncertainty, converge toward self-consistent values, and align more closely with each other within their intrinsic error margins. This underscores both the utility and limitations of each approach and highlights the importance of rigorous methods when quantitative accuracy is required in binding free energy predictions.

### Benchmark of Computational Binding Free Energies through Different Methods against Experimental Values

To directly evaluate the ability of the all-atom amber99sb-*disp* force field—using both FEP and PMF calculations—to reproduce the experimentally reported selectivity for different PARP inhibitors, we now compare the simulation-derived free energy differences with the experimental binding affinities reported for PARP1 and PARP2. Experimental IC_50_ values (i.e., the concentration of the ligand required to reduce the activity of the target protein by 50%) are converted into binding free energies according to the following relationship:

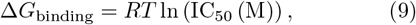

where *R* is the gas constant and *T* = 300 K. Experimental IC_50_ values are taken as follows: saruparib, 3 nM for PARP1 and 1400 nM for PARP2^70^; NMS-P118, 9 nM for PARP1 and 1390 nM for PARP2^15^; veliparib, 8.3 nM for PARP1 and 11 nM for PARP2^71^; and olaparib, 1.1 nM for PARP1 and 0.9 nM for PARP2^72^, and then converted into binding Δ*G* values (see Table S1 in SM for the experimental IC_50_ values used in this study, along with the chemical structures of the different PARP inhibitors). The associated uncertainty in the experimental binding free energies is estimated by propagating the standard deviation (*σ*) of the IC_50_ values reported across different assays in the ChEMBL database, providing an experimental error bar that reflects the variability inherent to heterogeneous biochemical measurements using the expression:

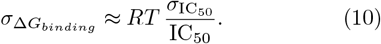

In Figure 3A, we present the comparison between experimentally derived differences in binding free energies between PARP1 and PARP2 (Δ(Δ*G*)) and those obtained from the MM/PBSA calculations. Consistent with the observations made earlier for the MM/PBSA *vs* FEP comparison, the overall agreement with experiment is weak: the correlation coefficient is low (*R*^2^ = 0.1269) and the deviation from an ideal linear relationship is significantly high (*D* = 7.54 kcal/mol, Eq. 8). A closer inspection reveals that this poor global correlation is primarily driven by veliparib, which appears as a clear outlier and does not follow the trend defined by the three other ligands. In contrast, the remaining three inhibitors—saruparib, NMS-P118, and olaparib—exhibit a more coherent linear trend between experimental and MM/PBSA-derived selectivities, mirroring the behavior observed in Figure 2B. This pattern reinforces that MM/PBSA struggles to capture subtle selectivity differences in highly alike ligand-protein complexes.

**FIG. 3.**
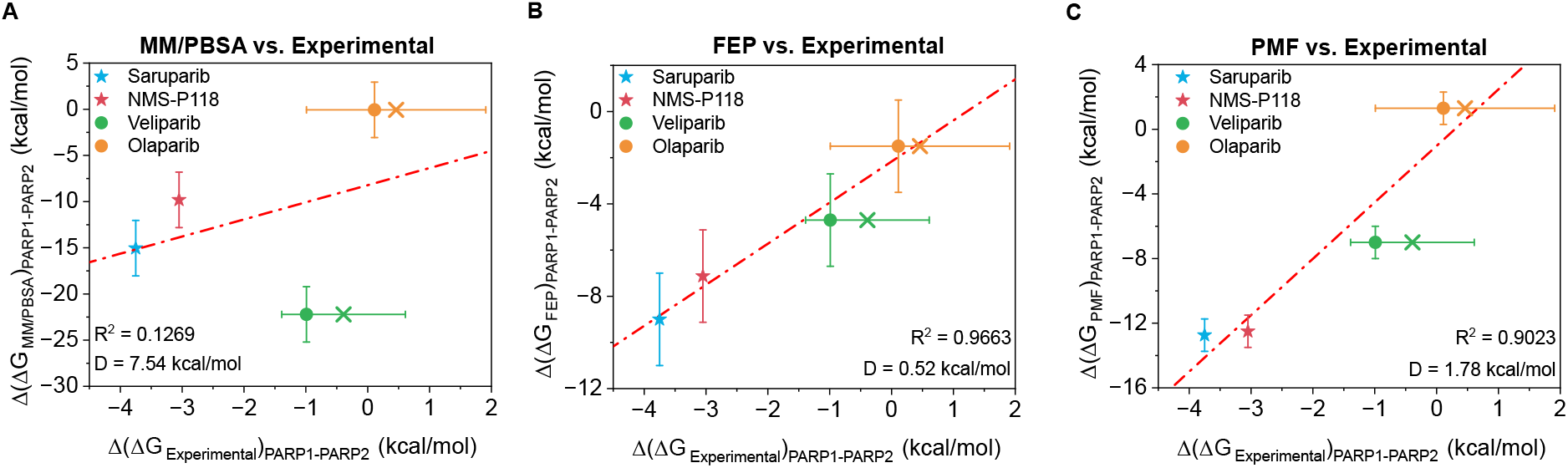
Comparison of the difference of binding free energy between PARP1 and PARP2 (Δ(Δ*G*)_*PARP* 1*−PARP* 2_) for four inhibitors: saruparib (blue star), NMS-P118 (red star), veliparib (green circle) and olaparib (yellow circle) using different computational approaches. (A) Correlation between MM/PBSA and experimental results^15,70–72^. (B) Correlation between FEP and experimental results^15,70–72^. (C) Correlation between PMF and experimental results^15,70–72^. The crosses indicate the mean IC_50_ values extracted from the ChEMBL database while filled circles depict the most cited IC_50_ value in literature. Error bars represent the standard deviation of the simulations and the estimated experimental uncertainty from different reported IC_50_ values using Eq. 10. Specific inhibitors of PARP1 are plotted as stars and non-specific inhibitors as circles. The red dashed line depict the linear regression of the data shown.

A notorious improvement is obtained with FEP, as shown in Figure 3B. Here, the correlation is remarkably strong, with *R*^2^ = 0.9663, demonstrating that FEP calculations in combination with the amber99sb-*disp*/OpenFF force fields capture near-quantitatively all of the experimental variance across the ligand set. The deviation from the ideal linear fit decreases significantly to *D* = 0.52 kcal/mol, well within the typical statistical uncertainty associated with FEP simulations. All four ligands fall close to the diagonal, and no significant outliers are observed, confirming that the force field accurately reproduces both the direction and magnitude of experimental selectivity shifts.

Finally, the PMF comparison is shown in Figure 3C. The PMF-derived differences in binding free energy between PARP1 and PARP2 also correlate strongly with experiments, yielding *R*^2^ = 0.9023 and a deviation of *D* = 1.78 kcal/mol. The magnitude of *D* lies close to the intrinsic uncertainty expected for PMF calculations (approximately 1 kcal/mol^26,63^). Importantly, the PMF approach correctly separates highly selective inhibitors such as saruparib and NMS-P118 from broadly active ones like veliparib and olaparib, mirroring experimental trends. A key advantage of PMF-based approaches is that, although positional restraints are required to define the reaction coordinate, the resulting free energy profiles preserve a physical interpretation^68^. Unlike FEP calculations, in which absolute binding free energies are not directly meaningful unless the thermodynamic loop is closed and only differences between carefully constructed alchemical states can be interpreted, a PMF describes the free energy landscape along a physically defined reaction coordinate (Fig. S1). This representation explicitly captures intermediate states, desolvation free energy barriers, and metastable configurations that arise during the ligand’s approach and accommodation within the binding pocket. Consequently, PMF calculations provide quantitative insight not only into the effective binding strength, but also on the mechanism, accessibility of the binding site, and the energetic cost associated with ligand insertion and rearrangement through a given dissociation pathway. As demonstrated in our previous work^68^, PMF profiles can yield quantitatively reliable relative binding free energies for closely related systems, while simultaneously offering mechanistic information—such as desolvation free energy barriers^69^—that are inherently inaccessible through standard alchemical FEP analyses based solely on Δ*G* differences.

It is also important to emphasize that the computational Δ*G* and Δ(Δ*G*) values are not expected to match experimental numbers quantitatively. Each method introduces intrinsic approximations that restrict the accessible conformational space of the protein–ligand complex. In particular, PMF simulations constrain the motion along a predefined reaction coordinate. These restraints reduce the accessible configurational entropy of the system and prevent exhaustive exploration of all relevant microstates, therefore limiting quantitative agreement with experiments beyond the intrinsic limitations of the employed force field. Even FEP, despite its rigorous statistical foundation, samples only a subset of the full conformational ensemble owing to the finite simulation time. Consequently, the appropriate comparison between computational and experimental values should preferably be based on relative trends and selectivity patterns rather than on absolute binding free energies.

Taken together, the results of Figure 3 reveal a clear hierarchy of predictive performance. MM/PBSA shows the weakest agreement with the experiment, exhibiting both the lowest correlation and the largest deviation values (Eq. 8). This reflects well–known limitations of endpoint approaches, which rely on single-configuration enthalpic estimates and do not explicitly account for the many microstates and entropic contributions that determine the binding free energy. On the other hand, PMF and FEP perform substantially better. PMF incorporates explicit solvent and partial configurational freedom along a reaction coordinate; and FEP achieves the most complete sampling of conformations and intermolecular interactions accessible during ligand binding within the protein pocket. As a result, both display stronger linear correlations with the experimental selectivities and significantly smaller deviation values. Among them, FEP yields the closest quantitative agreement, followed by PMF, which can be slightly affected by the positional restraints imposed during the reaction-coordinate sampling, as recently evidenced by us in Refs.^69,73^.

### Key Intermolecular Interactions Driving Ligand Selectivity in PARP1 Inhibition

To complement the free energy analysis presented above, we expand our dataset beyond the four PARP inhibitors originally examined by including four additional clinically relevant compounds. The motivation for this extension is twofold. First, incorporating a broader chemical diversity to assess how the trends in selectivity are determined by specific intermolecular contacts across the binding pocket. Second, this enlarged set enables a more reliable comparison between the two fastest computational approaches used to estimate binding affinity: MMGBSA and FEP. By analyzing the full panel of eight inhibitors with both methods, we aim to determine whether the additional compounds reinforce— or challenge—the selectivity patterns inferred from the initial dataset.

In Figure 4, we summarized the comparison between the computed and experimental difference in binding free energy between PARP1 and PARP2 (Δ(Δ*G*)_*PARP* 1*−PARP* 2_) for the full set of eight inhibitors. First, we report the MM/PBSA-derived selectivities (Figure 4A), while in Figure 4B we show the corresponding FEP predictions. The experimental IC_50_ values for the newly added inhibitors were taken as follows: rucaparib, 3.2 nM for PARP1 and 28.2 nM for PARP2^74^; niraparib, 3.8 nM for PARP1 and 2.1 nM for PARP2^75^; talazoparib, 2.59 nM for PARP1 and 0.89 nM for PARP2^76^ and pamiparib, 1.3 nM for PARP1 and 0.9 nM for PARP2^58^ (see Table S1 in SM for the experimental IC_50_ values used in this study, along with the chemical structures of the different PARP inhibitors). These values were subsequently converted into binding free energy differences (Δ*G*) (Eq. 9) for direct comparison with the computed results. When the four additional compounds are incorporated into the analysis, the overall performance of the two methods evolves in distinct ways.

**FIG. 4.**
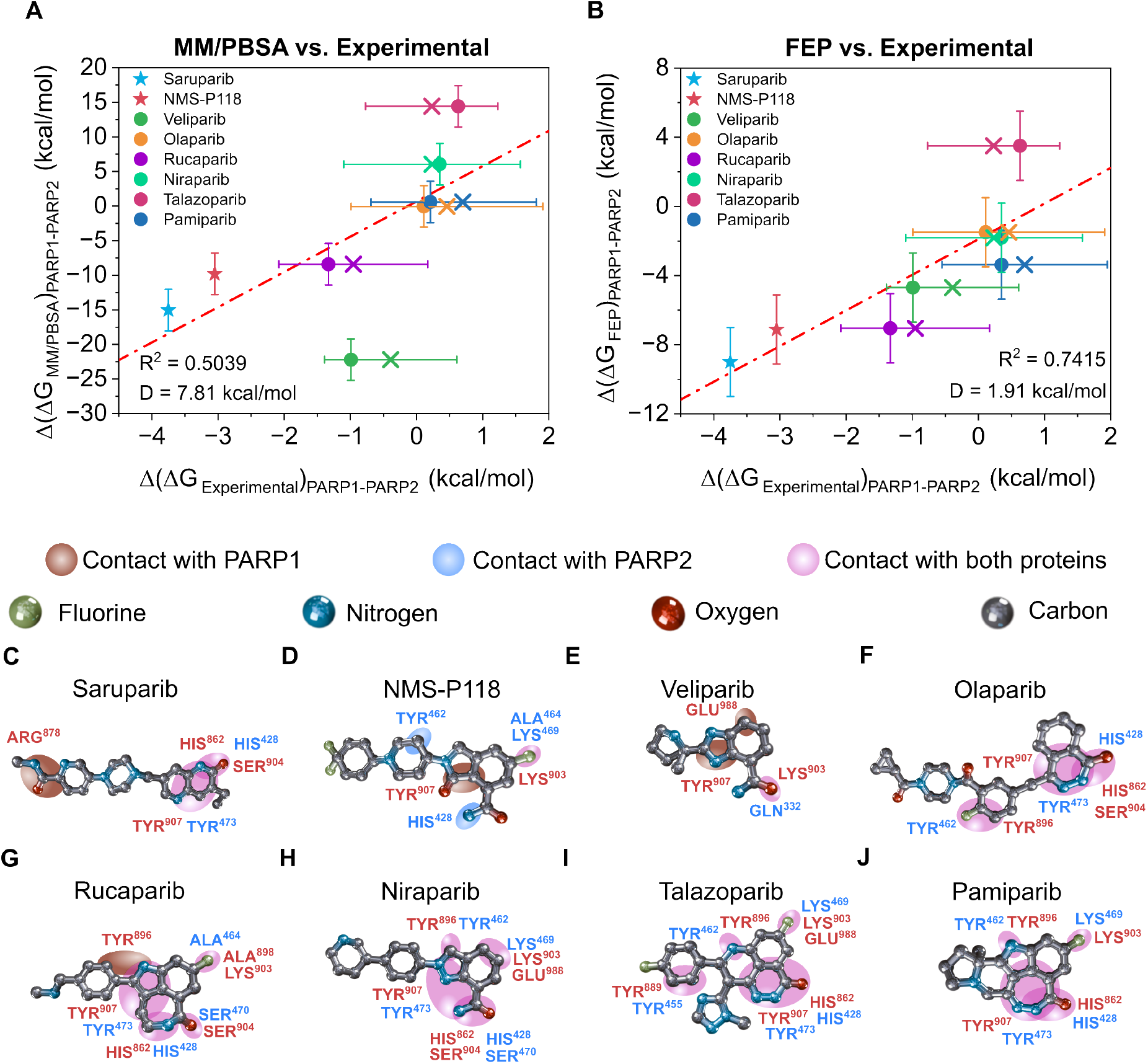
Comparison of binding free energy differences (ΔΔ*G*) between PARP1 and PARP2 for eight inhibitors: saruparib (blue star), NMS-P118 (red star), veliparib (green circle) and olaparib (yellow circle), rucaparib (purple circle), niraparib (mint circle), talazoparib (maroon circle) and pamiparib (dark blue circle) using different computational approaches. (A) Correlation between MM/PBSA and experimental results^15,58,70–72,74–76^. (B) Correlation between FEP and experimental results^15,58,70–72,74–76^. The crosses indicate the mean IC_50_ values extracted from the ChEMBL database while filled circles depict the most cited IC_50_ value from literature for each ligand. Error bars represent the standard deviation of the simulations and the estimated experimental uncertainty from different reported IC_50_ values using Eq. 10. Specific inhibitors of PARP1 are plotted as stars and non-specific inhibitors as circles. The red dashed line depict the linear regression of the data shown. Chemical structures of eight PARP inhibitors (C-J) annotated with their corresponding functional groups and regions establishing key contacts with the residues at the binding pockets in PARP1 (red labels) and PARP2 (blue labels). Panel C corresponds to saruparib, followed by NMS-P118 (D), veliparib (E), olaparib (F), rucaparib (G), niraparib (H), talazoparib (I), and pamiparib (J). Atom colors follow the scheme shown in the top legend: carbon (gray), nitrogen (blue), oxygen (red), and fluorine (green). Colored ellipses indicate protein–ligand contacts: brown marks regions contacting with PARP1, light blue marks regions contacting with PARP2, and purple denotes regions interacting with both proteins.

For MM/PBSA (Figure 4A), the agreement with experiment improves noticeably relative to the results obtained for the initial four inhibitors (Figure 3A). The coefficient of determination increases from *R*^2^ = 0.2673 to *R*^2^ = 0.5039, while the average deviation in binding energy difference remained roughly constant, *D* = 7.81 kcal/mol. This improvement in *R*^2^ indicates that the additional data dilutes the strong influence of the large error previously introduced by veliparib, which has disproportionately affected the regression when only four ligands are considered. As a result, the MM/PBSA yields a more defined correlation once the chemical diversity of the dataset is expanded. Moreover, if veliparib is excluded from the analysis, both statistical metrics improve substantially, with *R*^2^ increasing to 0.7898 and the deviation decreasing to *D* = 4.3 kcal/mol. This reduced deviation approaches the intrinsic error typically associated with MM/PBSA calculations^61^, highlighting that this method performs reasonably well for the remaining ligands when this potential outlier is removed. In contrast, FEP calculations (Figure 4B) show the opposite trend. Although it remains the more accurate method overall, its performance slightly deteriorates compared to the initial analysis (Figure 3B), giving *R*^2^ = 0.7415 and *D* = 1.91 kcal/mol. Nevertheless, the advantage of using FEP is still notable since its average deviation in binding free energy difference for the whole set of inhibitors remains over four times lower than that obtained through MM/PBSA calculations for the eight inhibitors.

To further test the robustness of our computational protocol, we perform additional FEP calculations for niraparib bound to two distant PARP-family members: PARP15 and TNK1, whose IC_50_ values (29200 nM^77^ and 40000 nM^78^, respectively) lie far outside the range observed for PARP1 and PARP2. As shown in Figure S2 of the SM, these FEP simulations also correctly reproduce the expected qualitative behavior, yielding markedly weaker binding for PARP15 and TNK1 relative to PARP1 and PARP2. This again confirms that our methodology is capable of resolving large selectivity differences across more divergent targets. By contrast, the selectivity between PARP1 and PARP2 represents an exceptionally demanding test case. The two catalytic domains are highly homologous, and the free energy differences that distinguish selective from non-selective inhibitors are rather small—often within the sub-kcal/mol regime where both experimental uncertainty and the intrinsic statistical noise of alchemical calculations become comparable to the signal itself. In this context, our FEP calculations performed here push the method to the edge of its practical resolution. The fact that FEP nonetheless recovers coherent and chemically meaningful trends across both the initial and expanded inhibitor sets underscores the robustness of the a99SB-*disp*/OpenFF allatom force field, as well as it supports the conclusion that the observed selectivity patterns are not artifacts of methodological choices since they are also consistent with previous PMF calculations (Fig. 2C).

It is also worth noting that Figure S3A–C in the SM presents the results of the MM/PBSA calculations without the entropic term (S3A), along with the corresponding MM/GBSA estimates without and with entropy (S3B and S3C, respectively). These data clearly show that omitting the entropic contribution systematically worsens the predictive performance in both approaches. More importantly, the comparison between MM/PBSA and MM/GBSA highlights a markedly superior performance of MM/PBSA for this class of systems: the PB-based solvation model yields more stable and consistent trends, whereas the GB variant exhibited a stronger sensitivity to both the dielectric environment and the conformational heterogeneity of the complexes^79^ (in Figure S4 of the SM are shown the charged amino acids of both protein active sites). This observation helps rationalize why, for structurally complex and highly charged binding pockets such as those examined here, MM/PBSA provides a more reliable framework than MM/GBSA for estimating relative binding free energies.

By extending the MD trajectories from FEP calculations to 0.2 *µs*, a detailed molecular analysis of residue-level contacts is also performed to explore whether structural and intermolecular interacting patterns can explain the selectivity differences observed between PARP1 and PARP2 (see Section VI of the SM for further details). While free energy calculations quantify the relative binding affinities, they do not explicitly reveal the microscopic molecular determinants responsible for protein selectivity. For this reason, we now examine the most frequent intermolecular contacts formed by each inhibitor within the catalytic binding site. The goal of this analysis is to identify recurring structural motifs—such as electrostatic interactions, conserved aromatic stacking, or differences in hydrogen-bonding networks—that might serve as key markers of molecular preference. By comparing the contact maps across selective and non-selective inhibitors, we sought to determine whether a consistent structural pattern, which correlates with experimentally known selectivity, may emerge.

For specific inhibitors, such as saruparib in Figure 4C, we reveal a binding mode in which key dominant contacts—HIS^862^ and the aromatic stacking with TYR^907^ (corresponding to the residues HIS^428^ and TYR^473^ in PARP2)—are partially shared between PARP1 and PARP2. The only two contacts that saruparib exclusively establishes in PARP1 are with ARG^878^ and SER^904^, which do not appear in PARP2 binding. Given that both proteins present an overall similar interaction pattern, the presence of these additional PARP1-specific interactions seems to be crucial in the experimentally observed selectivity for PARP1. In Figure 4D, NMS-P118 shows a different scenario. While both NMS-P118 and saruparib shared several stabilizing contacts—most notably TYR^907^ and LYS^903^—NMSP118 also forms a PARP1-exclusive interaction with TYR^907^, while PARP2 presents its own set of moderate stabilizing contacts with NMS-P118, including TYR^462^, HIS^428^, and ALA^464^ among others. This pattern suggests that cross-interactions with TYR^907^ and LYS^903^ in PARP1 might be critical for NMS-P118 behaving as a selective inhibitor of PARP1. Nevertheless, beyond the leading interactions with these two protein residues in PARP1, our data also suggests that selectivity likely arises from multiple, less dominant cooperative intermolecular contacts within each protein. The combination and synergy of these contacts further enhance non-shared interactions that differ significantly between PARP1 and PARP2 (Fig. S5-S8).

In contrast, poorly specific inhibitors, such as veliparib (Figure 4E), show an interaction pattern with both proteins that shares key contacts between heavy atoms of the ligand and protein residues including LYS^903^ in PARP1, or GLN^332^ in PARP2. Beyond these shared anchoring interactions, the remaining contacts are predominantly observed in PARP1, particularly with GLU^988^ and TYR^907^. This suggests that although veliparib engages both PARP1/PARP2 binding pockets through a similar recognition mechanism, the larger number of stabilizing interactions formed specifically with PARP1 may explain why both MM/PBSA and PMFs predict a slight preference toward PARP1. Olaparib (Figure 4F) represents the opposite scenario. All major contacts are shared between the two proteins, involving HIS^862^, TYR^896^, and TYR^907^ in PARP1, which correspond to HIS^428^, TYR^462^, and TYR^473^ in PARP2. The complete overlap in residue-level interactions further supported our FEP, PMF, and MM/PBSA calculations indicating that olaparib adopts essentially identical binding modes in both protein catalytic pockets. Such structural equivalence is fully consistent with the experimental observation that olaparib shows similar selectivity for both PARP1 and PARP2.

For rucaparib (Figure 4G), multiple intermolecular contacts are shared between PARP1 and PARP2, including TYR^907^, HIS^862^, SER^904^, and ALA^898^ in PARP1, as well as TYR^473^, HIS^428^, SER^470^, and ALA^898^ in PARP2. Notably, TYR^896^ and SER^904^ are observed exclusively in PARP1. The presence of these PARP1-specific interactions suggests that rucaparib may exhibit partial selectivity toward PARP1—as experimentally shown in Figure 4A—despite also forming multiple stabilizing contacts with PARP2. In contrast, niraparib (Figure 4H), talazoparib (Figure 4I), and pamiparib (Figure 4J) predominantly form contacts that are shared across both proteins. Key residues involved in these homologous interactions include TYR^896^, TYR^907^, HIS^862^, and LYS^903^, which are conserved in both PARP1 and PARP2 catalytic binding sites. The extensive overlap in the intermolecular contact patterns indicates that these inhibitors utilize a similar binding mechanism in both proteins and, consequently, cannot display significant selectivity. Overall, these structural observations reveal how contact connectivity controls ligand selectivity. Furthermore, our analysis provides valuable mechanistic and molecular insights into the chief residues that stabilize each ligand in both proteins (an energetic decomposition of the residues that contribute most strongly to ligand stabilization in each protein is shown in Figures S5–8), helping to elucidate potential chemical modifications across the ligand chemical space to enhance potency and specificity. Thus, structural contact analysis remains as an informative qualitative tool for interpreting free energy trends and guiding the development and optimization of nextgeneration oncologic drugs.

## CONCLUSIONS

In this work, we systematically evaluate the ability of three computational methods—MM/PBSA, FEP and PMF—to predict ligand inhibitor selectivity in two protein paralogues, PARP1 and PARP2, used as therapeutic targets in precision oncology. Our multi-method analysis reveals a clear progression in both methodological rigor and predictive performance. MM/PBSA, which relies on static energy-based estimations with implicit solvent, provides a rapid and qualitative view of relative selectivity. However, its predictions do not consider molecular conformational rearrangements, solvent-mediated interactions or real entropic contributions. Nevertheless, despite these aggressive approximations, MM/PBSA captured general trends among PARP1/PARP2 inhibitors (Fig. 4A), suggesting that it can serve as a useful initial screening tool for ligand selectivity.

FEP calculations using the all-atom a99SB-*disp* and OpenFF force fields, in contrast, achieve a much higher level of predictive accuracy. By explicitly sampling protein–ligand interactions and solvent dynamics, FEP calculations using this all-atom force field reproduce experimental selectivity patterns with remarkable fidelity. Across both the initial and expanded inhibitor sets, FEP consistently captures the trend and magnitude of binding free energy differences between PARP1 and PARP2, demonstrating that phase-space sampling is essential for resolving the subtle energetic distinctions that dictate selectivity between highly homologous proteins (Fig. 3B and Fig. 4B). Finally all-atom PMF calculations also using the a99SB-*disp* and OpenFF force fields show strong agreement with experimental data (Fig. 3C), correctly differentiating selective from non-selective inhibitors. Nevertheless, this method—which is the most computational expensive of the three studied techniques (Fig. 2A)—exhibits slight smoothing of energetic differences due to the required constraints of choosing a predefined dissociation pathway, which can partially limit discrimination among highly selective ligands.

Furthermore, our structural analyses elucidate the contact connectivity controlling ligand selectivity in PARP1. We reveal that while highly selective inhibitors such as saruparib and NMS-P118 form few protein-specific contacts—such as ARG^878^ and TYR^907^ by saruparib, or LYS^903^ and TYR^907^ by NMS-P118—most clinically used PARP inhibitors interact extensively with residues conserved across both PARP1 and PARP2 (Figs. 4C-J). These findings highlight why computer simulations including contact connectivity analyses and free energy calculations are vital to rationalize and propose new chemical modifications to enhance ligand specificity and potency. Taken together, our work highlights the critical role of rigorous calculations in computational drug design and establishes a robust framework to guide the development of selective ligands for next-generation cancer therapeutics.

## Data availability

We provide the relevant data in the repository (GitHub link for the repository) to facilitate reproducibility of our results. In the repository we also give the necessary code to run the simulations and accessible instructions to obtain our results.

## ACKNOWLEDGMENTS

A. F. acknowledges funding from the Ramon y Cajal fellowship (RYC2021-030937-I) and Spanish National Grant (PID2022-136919NA-C33). A. R. T. acknowledges funding from the European Union Horizon 2020 research and innovation programme (grant agreement 803326 to R.C.-G.). J. R. E. acknowledges funding from Emmanuel College, the University of Cambridge, the Ramon y Cajal fellowship (RYC2021-030937-I), the Spanish scientific plan and committee for research reference PID2022-136919NA-C33, and the European Research Council (ERC) under the European Union’s Horizon Europe research and innovation program (grant agreement no. 101160499). A.O., N.D.-V, C.P. and L.P.-H. acknowledge funding from CRIS Cancer Foundation (AOF.C01CRIS and AOF.M01CRIS). This work has also been performed using resources provided by Archer2 (https://www.archer2.ac.uk/) funded by EPSRC Tier-2 capital grant EP/P020259/e829. The authors also thankfully acknowledge RES computational resources provided by Mare Nostrum 5 through the activities 2024-3-0001 and 2025-1-0009.

