## Supplementary Material for "Benchmarking Free Energy Computational Methods for Revealing the Interactions Driving PARP1 Selective Inhibition"

Alejandro Feito

*Department of Physical Chemistry, Universidad Complutense de Madrid,*
*Av. Complutense s/n, Madrid 28040, Spain and*
*Instituto Pluridisciplinar, Universidad Complutense de Madrid,*
*P.º de Juan XXIII, 1, Moncloa - Aravaca, 28040 Madrid, Spain*

Natàlia DeMoya-Valenzuela, Cristian Privat, and Lucía Paniagua-Herranz

*Experimental Therapeutics Unit, Hospital Clínico San Carlos (HCSC),*
*Instituto de Investigación Sanitaria San Carlos (IdISSC), Madrid, Spain*

Andrés R. Tejedor

*Department of Physical Chemistry, Universidad Complutense de Madrid,*
*Av. Complutense s/n, Madrid 28040, Spain*
*Yusuf Hamied Department of Chemistry, University of Cambridge,*
*Lensfield Road, Cambridge CB2 1EW, UK and*
*Instituto Pluridisciplinar, Universidad Complutense de Madrid,*
*P.º de Juan XXIII, 1, Moncloa - Aravaca, 28040 Madrid, Spain*

Marco DelValle-Carrillo and Javier Oller-Iscar

*Department of Physical Chemistry, Universidad Complutense de Madrid,*
*Av. Complutense s/n, Madrid 28040, Spain*

Sara Cembellín

*Department of Organic Chemistry, Universidad Complutense de Madrid,*
*Av. Complutense s/n, Madrid 28040, Spain*

Adiran Garaizar

*Data Science, Bayer AG, Germany*

Alberto Ocana*

*Experimental Therapeutics Unit, Hospital Clínico San Carlos (HCSC),*
*Instituto de Investigación Sanitaria San Carlos (IdISSC), Madrid, Spain and*
*PhAsIca Biosciences S.L, Calle Velázquez, 27, 28001 Madrid, Spain*

Jorge R. Espinosa[†]

*Department of Physical Chemistry, Universidad Complutense de Madrid,*
*Av. Complutense s/n, Madrid 28040, Spain*
*Yusuf Hamied Department of Chemistry, University of Cambridge,*
*Lensfield Road, Cambridge CB2 1EW, UK*
*Instituto Pluridisciplinar, Universidad Complutense de Madrid,*
*P.º de Juan XXIII, 1, Moncloa - Aravaca, 28040 Madrid, Spain and*
*PhAsIca Biosciences S.L, Calle Velázquez, 27, 28001 Madrid, Spain*

(Dated: December 29, 2025)

*
†

### SI. DETAILS OF THE MM/PB(GB)SA METHOD

The protein structure was prepared using PDB2PQR [1] to ensure residue compatibility with AMBER force fields, and protonation states were assigned with PROPKA [2] at pH 7.0. The ligand was protonated with OpenBabel 3.1 [3] and parameterized with Antechamber [4] in AmberTools24, applying the GAFF [5] force field and AM1-BCC charge assignment. Missing parameters were generated with parmchk2. The protein–ligand complex was parameterized using tLeap, applying ff99SB-disp [6], GAFF, and TIP4P-*disp* [6] for the protein, ligand, and water molecules, respectively. The system was solvated in a rectangular box with at least 12.0 Å of solvent padding from any protein atom and 1.0 Å between any residue atom and the nearest solvent molecule to avoid steric clashes; $Na^+$ and $Cl^-$ ions were added to neutralize the system. Energy minimization was performed to relax the system after parameterization and solvation in three stages: (i) restraints on the entire complex, (ii) restraints on the protein backbone, and (iii) no restraints, each involving 2,000 steepest-descent and 3,000 conjugate-gradient steps with a 10 Å cutoff. The unrestrained stage was repeated three times (15,000 steps total), applying a restraint force constant of $50\ \text{kcal} \cdot \text{mol}^{-1} \cdot \text{Å}^{-2}$ when required.

MMPB/GBSA binding free energy calculations were performed for minimized structures. The solvent was stripped, and three topologies—complex, receptor, and ligand—were generated using mbondi2 radii. Binding free energies were computed using Poisson–Boltzmann (PB) and Generalized Born (igb = 5) models.

### SII. SYSTEM PREPARATION AND SIMULATION DETAILS

All-atom simulations were performed using the GROMACS simulation package (version 2023) [7]. Protein–ligand complexes were prepared to explore interaction potentials under physiological conditions, at a temperature of 300 K and an ionic strength of 150 mM NaCl. All simulations employed the a99SB-*disp* force field [6] for the protein, the TIP4P-*disp* water model, and ligand parameters derived using the OpenFF toolkit [8], ensuring a consistent and accurate description of protein–ligand interactions and solvation effects. Systems were solvated in explicit water, and $Na^+$ and $Cl^-$ ions were added to neutralize the total charge and achieve the target salt concentration.

### SIII. DETAILS OF THE FEP METHOD

For the FEP simulations, the protein–ligand complexes were solvated in a cubic box with dimensions of 8.5 × 8.5 × 8.5 nm at 300 K and 150 mM NaCl. Free-energy calculations employed 31 Coulomb ($\lambda_{\mathrm{coul}}$) and 31 van der Waals ($\lambda_{\mathrm{vdw}}$) coupling parameters to ensure smooth transitions between states. The specific values used for each window were:

Coulomb coupling parameters:

```
0.0, 0.125, 0.25, 0.375, 0.5, 0.625, 0.75, 0.875, 1.0, 1.000, 1.00, 1.000,
1.0, 1.00, 1.0, 1.00, 1.0, 1.00, 1.0, 1.00, 1.0, 1.00, 1.0, 1.00, 1.0,
1.00, 1.0, 1.00, 1.0, 1.00, 1.0
```

Van der Waals coupling parameters:

```
0.0, 0.000, 0.00, 0.000, 0.0, 0.000, 0.00, 0.000, 0.0, 0.025, 0.05, 0.075,
0.1, 0.15, 0.2, 0.25, 0.3, 0.35, 0.4, 0.45, 0.5, 0.55, 0.6, 0.65, 0.7,
0.75, 0.8, 0.85, 0.9, 0.95, 1.0
```

Each window was simulated for 2.5 ns. Intramolecular interactions of the ligand were scaled during the transformations, and soft-core potentials were applied with $\alpha = 0.5$, $\sigma = 0.3$, and a power of 1 to avoid singularities in van der Waals interactions. Neighbor lists were updated automatically, and FEP and dH/d$\lambda$ calculations were performed every 20 and 100 steps, respectively. The simulations used the a99SB-*disp* force field for the protein and ions, TIP4P-*disp* for water [6], and ligand parameters derived from the OpenFF toolkit [8]. Each $\lambda$ window was equilibrated and sampled to obtain accurate estimates of the free-energy differences, which were then combined to compute the relative binding free energies between ligands and protein.

The free-energy analysis was performed using the `AlchemicalAnalysis` package, employing the Multistate Bennett Acceptance Ratio (MBAR) method [9] to combine the results from all $\lambda$ windows and obtain rigorous estimates of the relative binding free energies.

### SIV. DETAILS OF THE PMF METHOD

For the PMF simulations, the configurations were solvated in a box of 8.5 × 8.5 × 15 nm at 300 K and 150 mM of NaCl. The center-of-mass (COM) distance between the protein and the ligand was controlled using a harmonic umbrella potential with a pulling force constant of 10000 kJ $mol^{-1}$ $nm^{-2}$. Approximately 60 umbrella sampling windows were defined, spaced every 0.025 nm within the range. For production runs, positional restraints of 1000 kJ $mol^{-1}$ $nm^{-2}$ (in directions perpendicular to the pulling axis) were applied to the amino acid heavy atoms of four residues of the protein with the highest frequency of contacts along the simulation with the ligand (to avoid rotations), and the heavy atoms of the drug were placed at the center of the molecule to allow freedom of movement of the regions that predominantly interact with the protein during the simulation. Each window was simulated for 10 ns. The Weighted Histogram Analysis Method (WHAM) [10], as implemented in GROMACS version 2023, was used to reconstruct the free energy profiles, and the initial 2000 ps of each simulation were excluded from WHAM analysis to ensure equilibration. The force field parameters and topology for all ligands were derived using the OpenFF toolkit [8], ensuring compatibility with the all-atom simulations for the protein sequence using the a99SB-*disp* for the protein and ions and TIP4P-*disp* for the water [6] force field.

### SV. SEQUENCE OF PARP1 AND PARP2 AND PDB OF THE STRUCTURED DOMAINS AND SEQUENCE OF PARP1

PARP1 Active Site:

```
NNADSVQAKVEMLDNLLDIEVAYSLLRGGSDDSSKDPIDVNYEKLKTDIKVVDRDSEEAEIIRKYVKNTHATTHN
AYDLEVIDIFKIEREGECQRYKPFKQLHNRRLLWHGSRTTNFAGILSQGLRIAPPEAPVTGYMFGKGIYFADMVS
KSANYCHTSQGDPIGLILLGEVALGNMYELKHASHISKLPKGKHSVKGLGKTTPDPSANISLDGVDVPLGTGISS
GVNDTSLLYNEYIVYDIAQVNLKYLLKLKFNFKTSLW
```

The following Protein Data Bank (PDB) codes were used for the atomistic simulations: PARP1 with saruparib (9ETQ [11]), with olaparib (7KK4 [12]), with veliparib (7KK6 [12]), with NMS-P118 (5A00 [13]), with niraparib (7KK5 [12]), with rucaparib (6VKK [14]), with pamiparib (7CMW [15]) and with talazoparib (7KK3 [12]).

PARP2 Active Site:

```
TQKELSEKIQLLEALGDIEIAIKLVKTELQSPEHPLDQHYRNLHCALRPLDHESYEFKVISQYLQSTHAPTHSDY
TMTLLDLFEVEKDGEKEAFREDLHNRMLLWHGSRMSNWVGILSHGLRIAPPEAPITGYMFGKGIYFADMSSKSAN
YCFASRLKNTGLLLLSEVALGQCNELLEANPKAEGLLQGKHSTKGLGKMAPSSAHFVTLNGSTVPLGPASDTGIL
NPDGYTLNYNEYIVYNPNQVRMRYLLKVQFNFLQLW
```

The following Protein Data Bank (PDB) codes were used for the atomistic simulations: PARP2 with olaparib (4TVJ [12]), with veliparib (3KJD [16]), with NMS-P118 (4ZZY [13])with niraparib (8HLQ [17]), with rucaparib (8HKO [17]), with pamiparib (8HKS [17]) and with talazoparib (4PJV [18]).

PARP15 Active Site:

`PEHWTDMNHQLFCMVQLEPGQSEYNTIKDKFTRTCSSYAIEKIERIQNAFLWQSYQVKKRQMDIKNDHKNNERLL`
`FHGTDADSVPYVNQHGFNRSCAGKNAVSYGKGTYFAVDASYSAKDTYSKPDSNGRKHMYVVRVLTGVFTKGRAGL`
`VTPPPKNPHNPTDLFDSVTNNTRSPKLFVVFFDNQAYPEYLITFTA`

The following Protein Data Bank (PDB) code was used for the atomistic simulations: PARP15 with niraparib (7F43 [19]).

TNK1 Active Site:

`KEIGINAYGHRHKLIKGVERLLGGQQGTNPYLTFHCVNQGTILLDLAPEDKEYQSVEEEMQSTIREHRDGGNAGG`
`IFNRYNVIRIQKVVNKKLRERFCHRQKEVSEENHNHHNERMLFHGSPFINAIIHKGFDERHAYIGGMFGAGIYFA`
`ENSSKSNQYVYGIGGGTGCPTHKDRSCYICHRQMLFCRVTLGKSFLQFSTMKMAHAPPGHHSVIGRPSVNGLAYA`
`EYVIYRGEQAYPEYLITYQIMKPEAPSQTATAAEQKT`

The following Protein Data Bank (PDB) code was used for the atomistic simulations: TNK1 with niraparib (7KKP [12]).

### SVI. CALCULATING CONTACT MAPS

Computation of intermolecular and intramolecular contact maps within protein condensates is performed from all-atom trajectories. In this analysis, molecular contacts are identified using a fixed distance criterion of 4 Å. Specifically, for each amino acid, the center of mass of its side chain is computed, and a contact is recorded whenever any heavy atom of the ligand lies within 4 Å of this center of mass. This definition provides a consistent geometrical measure of residue–ligand proximity that is independent of residue-specific excluded-volume parameters while still capturing the relevant short-range interactions.

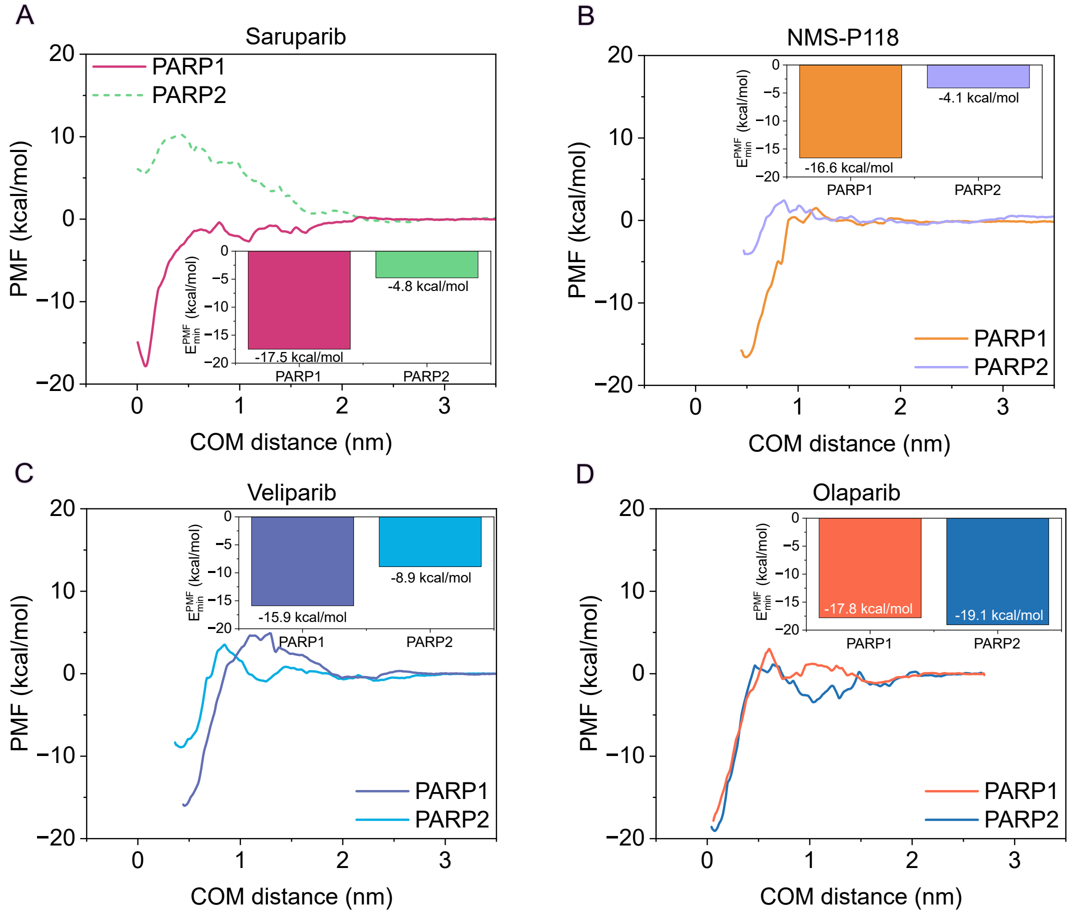


**FIG. S1:** Atomistic PMF dissociation profiles between each ligand and PARP1/PARP2 under physiological NaCl concentration (150 mM) and room conditions, in explicit solvent and ions, using the a99SB-disp force field. The center-of-mass (COM) distance between the protein and the ligand is used as the reaction coordinate. Curves are shown for saruparib (A), NMS-P118 (B), veliparib (C), and olaparib (D). Insets highlight the estimated minimum of the PMF for each complex, comparing PARP1 and PARP2. The dashed line of PARP2 with saruparib indicates that such configuration was generated through an MD trajectory rather than from equilibrated structurally resolved PDBs.

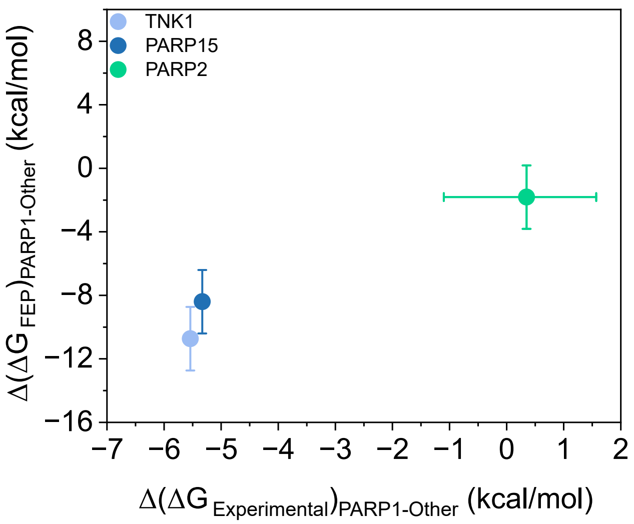


**FIG. S2:** Comparison between experimental and FEP-predicted PARP1 selectivity for three off-targets (TNK1, PARP15, and PARP2). The x-axis reports the experimental relative selectivity, $\Delta(\Delta G_{\text{Experimental}})$PARP1–Other, while the y-axis shows the corresponding FEP-predicted value, $\Delta(\Delta G\text{FEP})_{\text{PARP1–Other}}$.

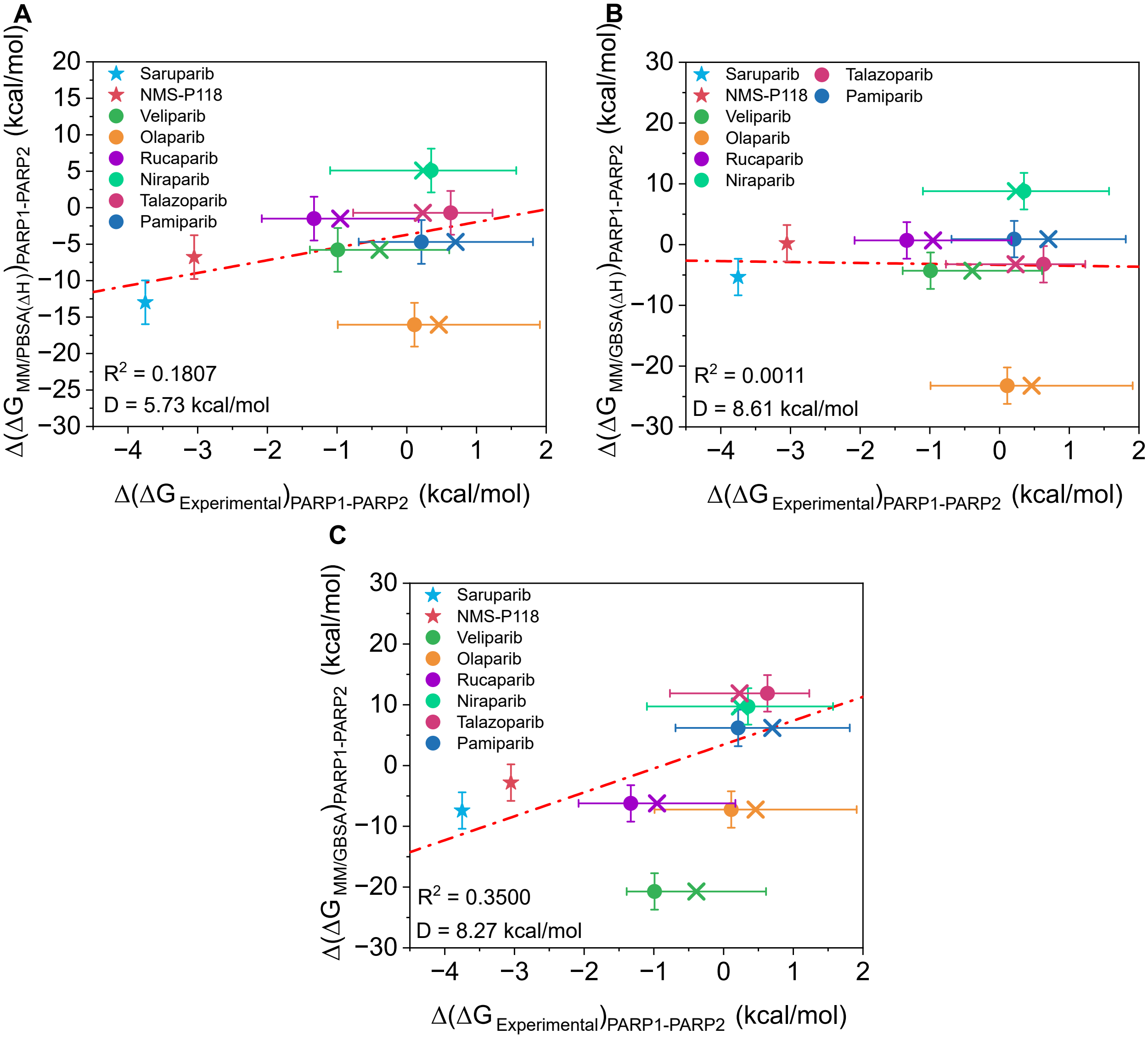


**FIG. S3:** Comparison of binding free-energy differences ($\Delta\Delta G$) between PARP1 and PARP2 for eight inhibitors: saruparib (blue star), NMS-P118 (red star), veliparib (green circle) and olaparib (yellow circle), rucaparib (purple circle), niraparib (mint circle), talazoparib (maroon circle) and pamiparib (dark blue circle) using different computational approaches. (A) Correlation between MM/PBSA method (only the enthalpic term) and experimental results [13, 15, 20–25]. (B) Correlation between MM/GBSA (only the enthalpic term) and experimental results [13, 15, 20–25]. (C) Correlation between MM/GBSA and experimental results [13, 15, 20–25]. The crosses indicate the mean $IC_{50}$ values extracted from the ChEMBL database. Error bars represent the standard deviation of the simulations and the estimated experimental uncertainty from different reported $IC_{50}$ values using Eq. 12. Specific inhibitors of PARP1 are plotted in stars and non-specific inhibitors in circles. The red dashed line depict the linear regression.

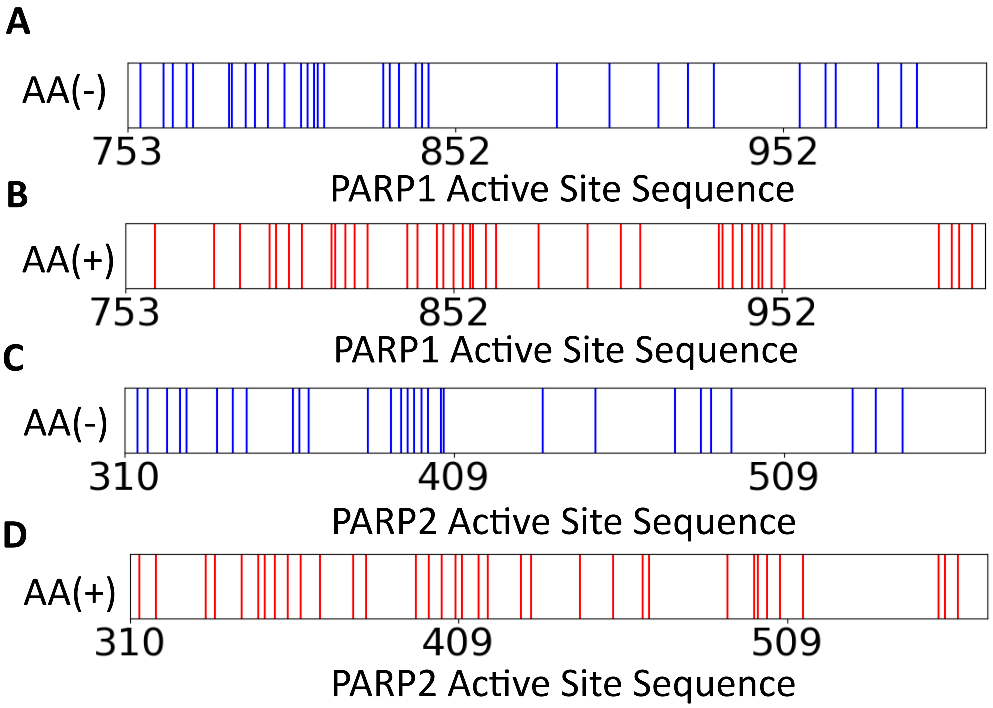


**FIG. S4:** Schematic representation of charged amino acids in the active sites of PARP1 and PARP2. (A) Negatively charged amino acids (AA(-), blue) in the PARP1 active site. (B) Positively charged amino acids (AA(+), red) in the PARP1 active site. (C) Negatively charged amino acids (AA(-), blue) in the PARP2 active site. (D) Positively charged amino acids (AA(+), red) in the PARP2 active site.

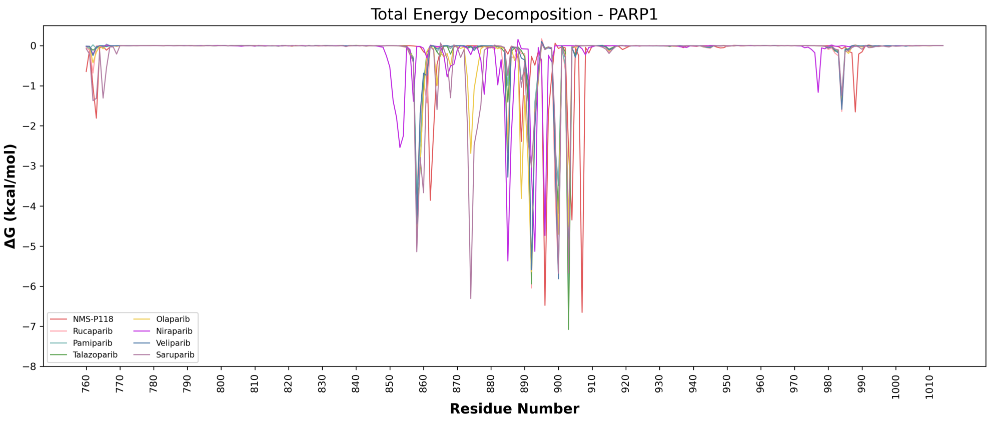


**FIG. S5:** Total energy decomposition of the different ligands in this study for the active site of PARP1.

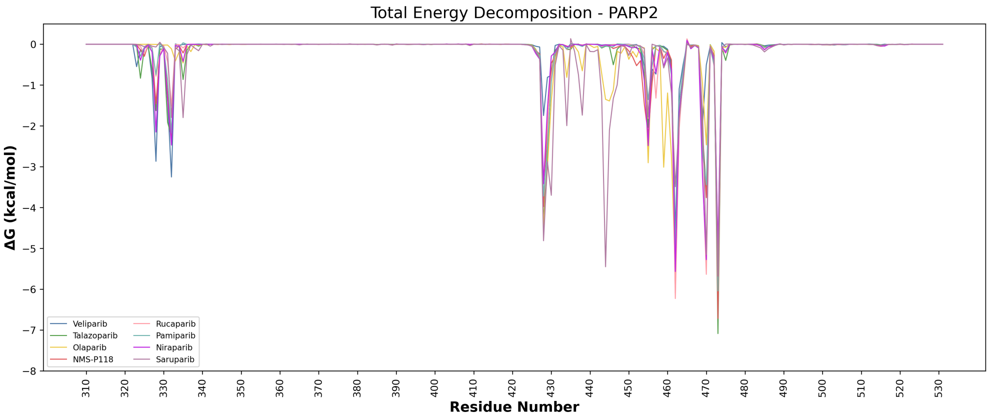


**FIG. S6:** Total energy decomposition of the different ligands in this study for the active site of PARP2.

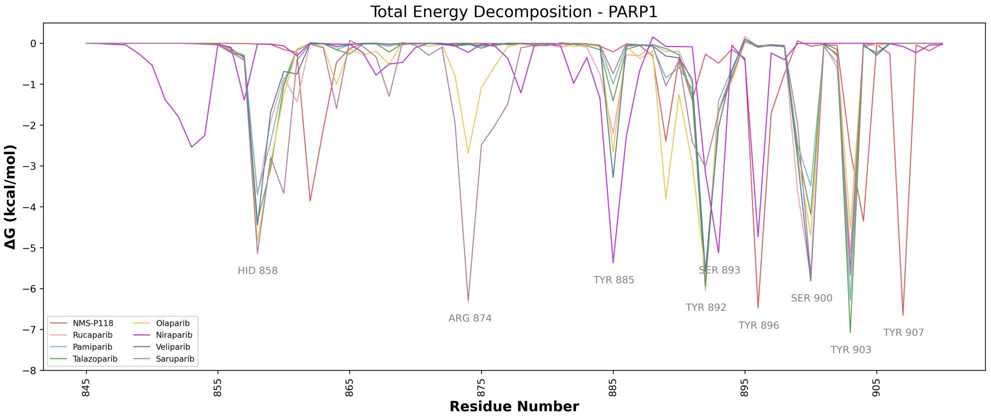


**FIG. S7:** Total energy decomposition of the different ligands in this study for the pocket of the active site of PARP1.

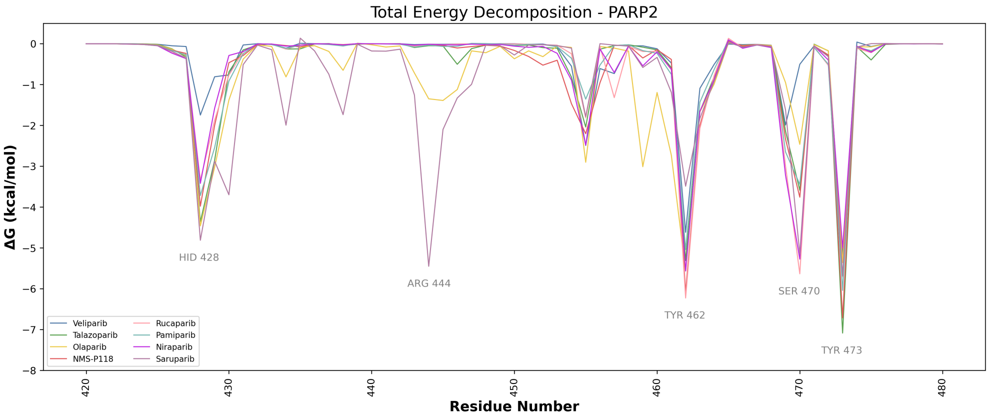


**FIG. S8:** Total energy decomposition of the different ligands in this study for the pocket of the active site of PARP2.

**TABLE S1:** Experimental $IC_{50}$ values for PARP inhibitors used in this study.

| Compound | Structure | $IC_{50}$ PARP1 (nM) | $IC_{50}$ PARP2 (nM) |
|---|---|---|---|
| Saruparib | 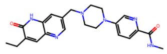 | 3.0 [20] | 1400 [20] |
| NMS-P118 | 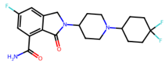 | 9.0 [13] | 1390 [13] |
| Veliparib | 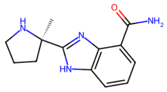 | 8.3 [21] | 11.0 [21] |
| Olaparib | 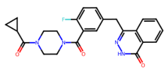 | 1.1 [22] | 0.9 [22] |
| Rucaparib | 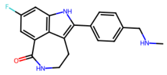 | 3.2 [23] | 28.2 [23] |
| Niraparib | 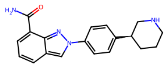 | 3.8 [24] | 2.1 [24] |
| Talazoparib | 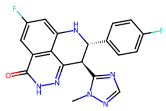 | 2.59 [25] | 0.89 [25] |
| Pamiparib | 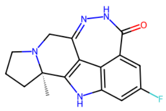 | 1.3 [15] | 0.9 [15] |
